# CombPDX: a unified statistical framework for evaluating drug synergism in patient-derived xenografts

**DOI:** 10.1101/2021.10.19.464994

**Authors:** Licai Huang, Jing Wang, Bingliang Fang, Funda Meric-Bernstam, Jack A. Roth, Min Jin Ha

## Abstract

**Motivation:** Anticancer combination therapy has been developed to increase efficacy by enhancing synergy. Patient-derived xenografts (PDXs) have emerged as reliable preclinical models to develop effective treatments in translational cancer research. However, in most PDX combination experiments, PDXs are tested on single dose levels and dose-response surface methods are not applicable for testing synergism.

**Results:** We propose a comprehensive statistical framework to assess joint action of drug combinations from PDX tumor growth curve data. We provide various metrics and robust statistical inference procedures that locally (at a fixed time) and globally (across time) access combination effects under classical drug interaction models. Integrating genomic and pharmacological profiles in non-small-cell lung cancer (NSCLC), we have shown the utilities of combPDX in discovering effective therapeutic combinations and relevant biological mechanisms.

**Availability:** We provide an interactive web server, combPDX (https://licaih.shinyapps.io/CombPDX/), to analyze PDX tumor growth curve data and perform power analyses.

**Supplementary information:** Supplementary data are available at *Bioinformatics* online.

## 1 Introduction

Combinations of anticancer drugs have been developed to circumvent mechanisms of resistance to yield clinical benefit and lower toxicity (Dawson and Carragher, 2014; Wright, 2016). Recently, in vitro high-throughput combinatorial screening data have been enabling the assessment of large number of drug combinations at various dose levels (Mathews Griner, et al., 2014; Narayan, et al., 2020; Zagidullin, et al., 2019). In these experiments, a pair of drugs are plated in a dose-response matrix block and the data at various combinations of dose levels are analyzed to quantify the degree of combination effects. The joint effects are categorized into synergistic, additive, and antagonistic, which imply enhanced, independent, and reduced effect, respectively, when two drugs are present together. The responses obtained from multiple combinations of dose levels are compared against the expected response under null models where no combination effect is present. The classical reference null models include Highest single agent (HSA) (Berenbaum, 1989; Geary, 2013; Lehár, et al., 2007), Loewe additivity (Frei, 1913; Loewe, 1928; Loewe, 1953; Loewe and Muischnek, 1926) and Bliss independence (BI) (Berenbaum, 1989; Bliss, 1939; Geary, 2013; Greco, et al., 1995). Recent application developments, such as DrugComb (Zagidullin, et al., 2019), SynergyFinder (Ianevski, et al., 2017), and Combenefit (Di Veroli, et al., 2016) provide computational tools to analyze drug combination screening data based on these reference models.

Patient-derived xenografts (PDXs) have emerged as reliable preclinical models to develop new treatments and biomarkers in translational cancer research (Jung, et al., 2018). The PDX models are developed by implanting tumors from patients into mice; this method has been suggested to more accurately reflect clinical outcomes (Gao, et al., 2015; Izumchenko, et al., 2017). These successes have led to a rapid accumulation and availability of large-scale PDX collections for drug discovery in cancer (Conte, et al., 2019; Doroshow, 2016; Gao, et al., 2015). In the PDX experiments to evaluate drug efficacy, tumor volumes of each individual mouse are measured at the initiation of the study and periodically throughout the study. This usually continues until the tumor volume reaches a certain value, resulting in incomplete longitudinal tumor volume data. Due to the high cost of in vivo studies in animals, a common combination experiment for a PDX model includes four treatment groups, control (C), two monotherapies (A and B), and combination therapy (AB) with fixed doses to minimize the number of animals required per group. In the fixed dose experiment, the dose-response surface methods are not applicable, and the joint action of drugs should be evaluated at a fixed dose combination.

The statistical framework that assesses the joint action of drug combinations with fixed doses is not well developed. Wu, et al. (2013; 2012) proposed interaction indices and the statistical inferential procedure based on surviving fraction of cells and survival endpoints. Demidenko and Miller (2019) proposed a log-linear model on the tumor volumes by assuming that they follow exponential growth. However, these are limited to the Bliss independence model. A distinctive set of mathematical definitions might lead to different quantifications of the degree of joint action.

In this article, we propose a comprehensive statistical framework to calculate combination indices and the inferential procedures that are robust to potential outliers and errors in PDX experiments. We considered three most well-established reference models: HSA, response additivity (RA), and BI in a unified statistical quantification method. We present a user-friendly web server, combPDX, to compile the joint actions over time from PDX tumor growth data and to provide a power analysis tool to facilitate designs of PDX combination studies. Applying our methods to non-small-cell lung cancer (NSCLC), we show the utilities of our framework in finding underlying mechanisms of combination drug action using gene expression profiles for PDX models.

## 2 Methods

### 2.1 Overview

The pipeline of combPDX includes three steps to assess combination effects as well as the power analysis procedure (Figure 1). Longitudinal raw tumor volume measurements are collected from four treatment groups (C, A, B and AB) and the tumor growth curves are displayed (Figure 1a). For each individual mouse, the response at each time point is determined by computing the relative tumor volume to adjust for heterogeneous initial tumor measurements across animals, and the missing relative tumor volumes at time t are interpolated using the neighboring measurements (Figure 1b). At each time point, we determine the treatment effects of A, B, and AB compared to the control group C (Figure 1c). Based on the treatment effects, we provide the combination indices under HSA, RA, and BI, and the corresponding 95% confidence intervals (Figure 1d). We finally implement a power analyses tool under the three reference models (Figure 1e).

**Figure 1.**
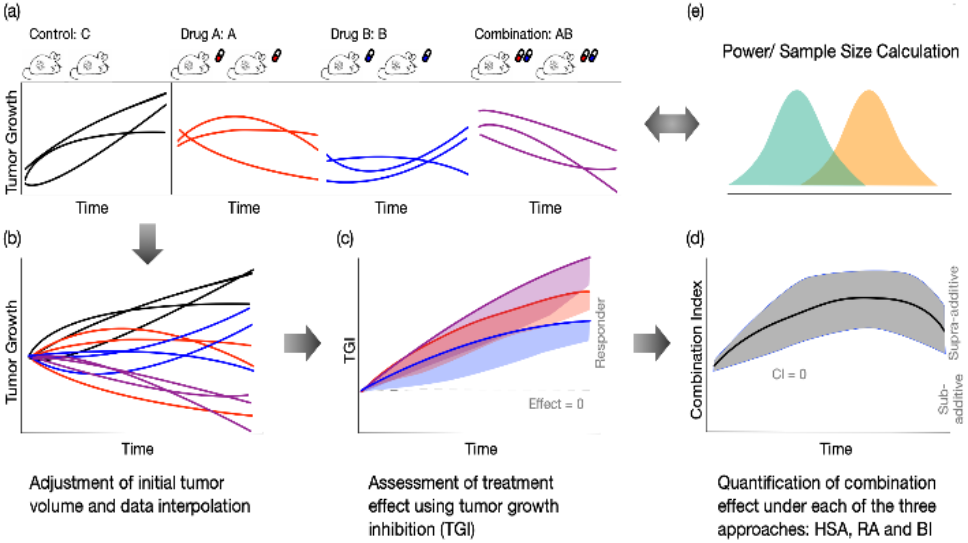
Overview of the analysis pipeline of combPDX. (a) Tumor volume measurements from four treatment arms (C, A, B, and AB) are collected from PDX experiments. (b) For an individual mouse, the response at each time point is determined by computing relative tumor volume to adjust for heterogeneous initial tumor measurements. (c) The drug effect for each treatment group (A, B, and AB) relative to the control group is quantified by tumor growth inhibition (TGI). (d) The combination effect under each reference model (HSA, RA, and BI) is assessed using a combination index. In addition, (e) sample size calculation and power analysis are provided.

### 2.2 Tumor Volume Data Processing

The combPDX requires a long data matrix as an input where each of the tumor volume measures are stacked by rows with four columns: mice ID, treatment, day, and tumor volume. Table S1 presents the description of these metrics and Table S2 shows an example of input data. Due to the variation of initial tumor volumes across mice, the response for an individual animal at each time point is defined by the *relative tumor volume*, which is the raw tumor volume divided by the initial tumor volume of the mouse. We denote *v*_*t*_ as the relative tumor volume for a mouse at time *t* (Section S1.1). For subjects with no tumor volume measurement at time *t*, but with flanking volume measurements at time *t*_0_ and *t*_1_, we use linear interpolation to impute the relative tumor volume at time *t*:

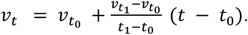

Denote the relative tumor volume for an individual mouse in group *g* = *C*, *A*, *B*, *AB* as *v*_*g*_, which follows an independent identical distribution with mean *μ*_*g*_ and variance 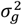. The mean and variance parameters *μ*_*g*_ and 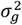 can be estimated by the sample mean 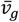 and sample variance 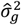. Our analytic inference procedure in this paper is based on the central limit theorem, 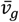 follows normal distribution with mean *μ*_*g*_ and variance 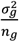, where *n*_*g*_ is the sample size in treatment group *g* (Section S1.1).

Combined with Boostrap procedures, we re-calibrate the null distribution of test statistics to propose a robust statistical framework to any deviations from the theoretical distribution.

### 2.3 Determination of Treatment Effect

To access the combination effect of two drugs, we take effect-based approaches that directly compare the effect of the combination to the effects of its individual components. The effect of a treatment group (A, B or AB) is defined by the antitumor activity compared with the control group (C). At a given time point, treatment effect for a group *g* is quantified by the mean reduction in the relative tumor volumes between treatment and control groups divided by the control mean

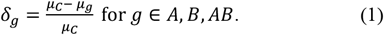

A large *δ*_*g*_ value indicates a strong treatment effect. For combination experiments, we expect that the mean relative tumor volumes of the treatment groups *μ*_*A*_, *μ*_*B*_ and *μ*_*AB*_ are less than the control tumor volume *μ*_*C*_, which results in *δ*_*A*_, *δ*_*B*_ and *δ*_*AB*_ located between 0 and 1. While this has been widely used as the tumor growth inhibition (TGI) with predetermined cutoffs of declaring an antitumor activity (Houghton, et al., 2007; Mer, et al., 2019; Ortmann, et al., 2020), we incorporate statistical inferential procedures by constructing 95% confidence intervals. The lower bound of a one-sided *100(1-α*)% confidence interval for a combination index can be calculated using the Delta method,

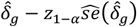

where 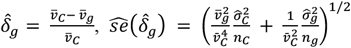, and *z*_1−*α*_ is the (1 − *α*)^th^ quantile of standard normal distribution (Section S1.2).

### 2.4 Combination Index (CI)

Based on the treatment effects *δ*_*g*_ evaluated for all three treatment groups A, B and AB, we aim to access the superiority of using drug combination AB to individual drugs A and B. Although there is no consensus on defining the synergistic action of two drugs (Roell, et al., 2017), we derive combination indices under the three popular reference models: (1) HSA (*CI*_*HSA*_), (2) RA (*CI*_*RA*_), and (3) BI (*CI*_*BI*_). All the CIs under the three models are calibrated to have the same implication: *CI* < 0, = 0, > 0 represent antagonistic, independent, and synergistic effects, respectively.

**Highest Single Agent (HSA)** (Berenbaum, 1989; Foucquier and Guedj, 2015; Geary, 2013; Lehár, et al., 2007) shows that the synergistic combination effect occurs when the combined effect is greater than the more effective individual component: 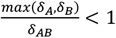. Under the HSA, we derive *CI* with respect to group means:

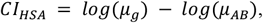

where *g* is chosen from *A* or *B* that has larger effect *δ*. If the two single agent effects are equal, *δ*_*A*_ = *δ*_*B*_, we choose the one that has the narrower confidence interval evaluated in Section 2.3. The *CI*_*HSA*_ is independent of the control experiment.

**Response Additive (RA)** (Foucquier and Guedj, 2015; Slinker, 1998) assumes that the fixed-dose two-drug combination has a linear additive effect under independence. A combination drug is considered synergistic if it shows a more enhanced effect than the sum of the two monotherapies’ effects: 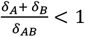. The corresponding *CI* can be derived with respect to group means as:

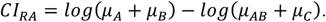

The **Bliss Independence (BI)** approach (Berenbaum, 1989; Bliss, 1939; Foucquier and Guedj, 2015; Geary, 2013; Greco, et al., 1995) is based the definition of independence on its probabilistic interpretation. Two drugs are independent if one drug’s presence does not affect the probability of another drug’s effect on tumor growth decay. Wu et al (2012) proposed an interaction index under such a definition using relative tumor volume. Assume that the treatment effects *δ*_*g*_ are the outcomes of a probabilistic process such that 0 ≤ *δ*_*g*_ ≤ 1. Note that *δ*_*g*_ in equation (1) takes a value between 0 and 1 if *μ*_*C*_ > *μ*_*g*_ for *g* = *A*, *B*, *AB*. Under BI, if the combination therapy is considered synergistic, we have 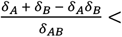 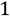. The combination index is given by

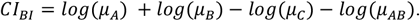

### 2.5 Calibration of confidence intervals for CIs

We develop statistical inferential procedures by deriving confidence intervals for the *CIs* under the asymptotic normality in addition to the bootstrap method. Using the Delta method, the standard errors of the indices are approximated by:

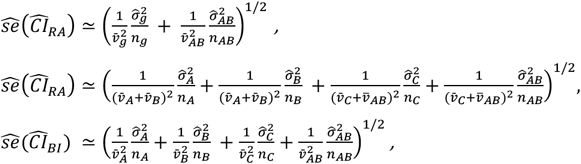

A two-sided 100(1 − *α*)% confidence interval is constructed as 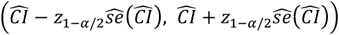 (see Derivations in Section S1.3-1.5).

The standard intervals based on asymptotic approximation to normal distributions can be inaccurate in practice due to skewness and heavier tails of tumor volume data. A robust bootstrap procedure is provided to construct confidence intervals (Efron and Tibshirani, 1994). Bootstrap is a resampling method that samples the original data with replacements iteratively to estimate test statistics or null distributions. To calculate bootstrap-t interval at a given time point, we repeat the following steps B times. For a given reference model, HSA, RA or BI, at the iteration b, we

1. Sample a size of *n*_*g*_ animals with replacement within each treatment arm *g*, and the corresponding tumor volume measurement for an animal is denoted by 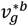.
2. Calculate combination index 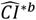 based on the bootstrap samples from the step 1, 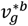 for *g* = *C*, *A*, *B*, *AB*.
3. Compute 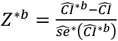 where 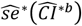 is calculated by the theoretical standard error calibrated in Section 2.5.

The *α*^*th*^ percentile of *Z*^\**b*^ is estimated by the value 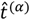 such that 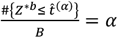. Finally, the bootstrap-t confidence interval is constructed by 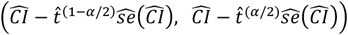. This bootstrap procedure adjusts for derivations from the asymptotic distribution of *CI* by recalibrating the percentiles.

### 2.6 Global Assessment of Combination Effect

The *CI* values assess combination effects at a given time point. We extend the procedure to a global assessment for any given study intervals of interest. Given reference model, the global *CI* is defined as the average of the *CI*s within the study interval of interest for time points 1, …, *T*:

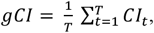

where *CI*_*t*_ is the combination index at time *t*. Due to correlations of tumor volume measurements within individual mouse, we conduct nested bootstrap procedure to construct a confidence interval for *gCI* without analytically specifying the variance *var*(*gCI*) (Efron and Tibshirani, 1994). The algorithm to construct confidence interval for *gCI* is similar to that in Section 2.5, with an additional nested layer to estimate 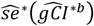. At the iteration b, we

1. Sample a size of *n*_*g*_ animals with replacement within each treatment arm *g*, the corresponding growth curve data for an animal are denoted by 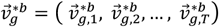.
2. Calculate combination index 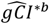 based on the 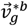, *g* = *C*, *A*, *B*, *AB*. We repeat the following nested bootstrap procedure *L* = 25 times to estimate 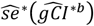,

2.a. Sample a size of *n*_*g*_ growth curves with replacement from 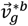 within each group and denote the corresponding growth curve data as 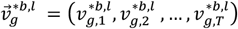 for each animal.
2.b. Calculate combination index 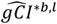 based on 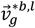, *g* = *C*, *A*, *B*, *AB* The standard error of each resampled data set can be estimated by

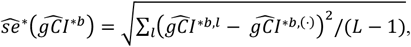

where 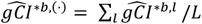.
3. Compute 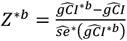.

The α^th^ percentile of *Z*^\**b*^ is estimated by the value 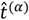 such that 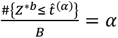. Finally, the bootstrap-t confidence interval is 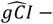 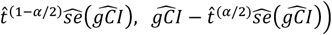.

### 2.7 Power Analysis

Based on the CIs under the HSA, BI and RA reference models, we provide power analysis tool to design PDX combination experiments. Under the null hypothesis of independent action of a drug combination, *H*_0_: *CI* = 0, where the *CI* follows asymptotic normal distribution *N*(0, *var*(*CI*)). Assume that under the alternative hypothesis *H*_*A*_: *CI* = *γ*, where *CI* follows normal distribution *N*(*γ*, *var*(*CI*)). Therefore, with prespecified values *μ*_*g*_, *σ*_*g*_, *n*_*g*_ for each group *g*, the power is calculated as

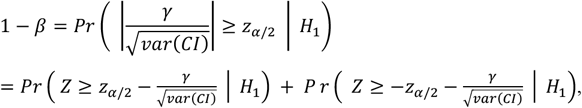

where *α*, *β* are the desired Type I and Type II error rates.

Given mean tumor volumes, Figure 2a illustrates the minimum *δ*_*AB*_ having synergistic effect under each reference model given *δ*_*A*_ and *δ*_*B*_, and we also mathematically compared *CI*_*HSA*_, *CI*_*RA*_ and *CI*_*BI*_ in Section S1.6. Given mean tumor volumes, we have 0 ≤ *CI*_*RA*_ ≤ *CI*_*BI*_ ≤ *CI*_*HSA*_, which implies that HSA model provides the most relaxed procedure while RA is the most conservative. Combined with the statistical inference procedure using the asymptotic normality, the power curves, Figure 2b and 2c for sample sizes of 5 and 10, respectively indicate that RA (HSA) require the largest (smallest) effect sizes of the combination to achieve the same statistical power when the effect sizes of monotherapies and standard deviations are fixed.

**Figure 2:**
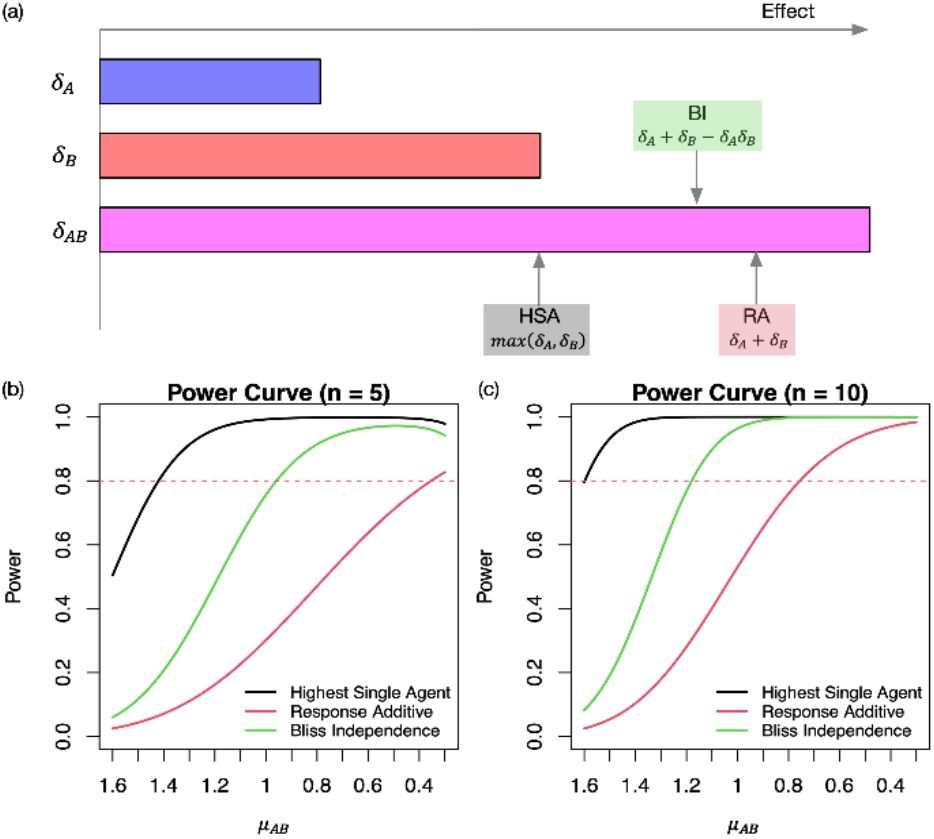
Comparison of HSA, BI and RA. models under the hypothetical scenario where *μ*_*C*_ = 2.8, *μ*_*A*_ = 2.4 *and μ*_*B*_ = 2. (a) Given *δ*_*A*_ *and δ*_*B*_, the arrows indicate the minimum *δ*_*AB*_ values that have synergistic effects (*CI* > 0). (b)-(c) Statistical power varies by the mean tumor volume for the combination AB when the sample size is 5 and 10 at the 0.05 significance level. The standard errors of relative tumor volumes are set to 0.7 for the control group C and 0.3 for the treatment groups A, B and AB

## 3 Results

### 3.1 Simulation Studies

We conducted a series of simulations to examine the performance of the proposed approaches. The expected tumor volumes under singe-agent treatment or control group were generated by the Gompertz tumor growth model over time (Ribba, et al., 2014; Winsor, 1932). Then the expected tumor volumes for combination drug were generated under each of the reference model (HSA, RA, and BI). The expected tumor growth curves for these simulation settings are shown in Figure S2. For each setting, we generated 1000 replicate datasets with sample sizes of 5 or 10 per group. The detailed data generating process is described in Section S1.7. We evaluated the CIs with the confidence intervals at day 21 obtained from all the three reference models under each of the simulation scenarios. We compared the inferential methods with/without the bootstrap procedures based on 1000 replications. The coverage probabilities are summarized in Table S3. When sample size is 5 in each group, the confidence intervals without the bootstrap procedure were slightly narrower than nominal level, resulting in inflated type I error under the null hypothesis. When we have more sample size (n=10) in each group, confidence intervals without bootstrap become close to the nominal coverage probability, and the confidence intervals with and without bootstrap tend to agree on each other. Overall, the bootstrap procedure helps in constructing more accurate confidence intervals when small sample size while both procedures provide valid statistical inferential performance when sample size becomes larger.

### 3.2 Evaluation of drug combinations of KRT232, navitoclax, and trametinib in NSCLC

We performed the analysis of a real study of the antitumor activity of the combination of KRT232, navitoclax, and trametinib using the PDX models for NSCLC, where a total of 28 PDX models were tested (Chen, et al., 2019; Hao, et al., 2015; Zhang, et al., 2020). For each PDX tumor model, mice were randomized to the four treatment arms, and the tumor volume for each mouse was recorded every 2-3 days. 20 combination therapy experiments having sufficient samples sizes were selected (Section S2). The treatment information for the selected experiments is summarized in Table S4. Table S5 summarizes the resulting *CIs* at day 10 and *gCI*s under the three reference models for each PDX model. trametinib and navitoclax combination shows the synergism and figure 3 displays the output from our combPDX analysis. Raw tumor volumes for all mice across the four groups are displayed in panel (a). After the preprocessing step in Section 2.2, the average relative tumor volumes with 1-sd bars are displayed in panel (b). Then the effect sizes of each treatment groups A, B, AB relative to the control group are determined with 95% confidence intervals in panel (c) from section 2.3, implying that all treatments trametinib, navitoclax, and the combination treatments have the significant antitumor activity compared to the control in the PDX experiment. Trametinib and navitoclax combination has the synergistic effect under HSA and BI from day 6 to day 18 and from day 8 to day 11, respectively (Figure 3d-f and Table S5). This combination is currently under clinical investigation for the treatment of NSCLC (Kim and Giaccone, 2018).

**Figure 3:**
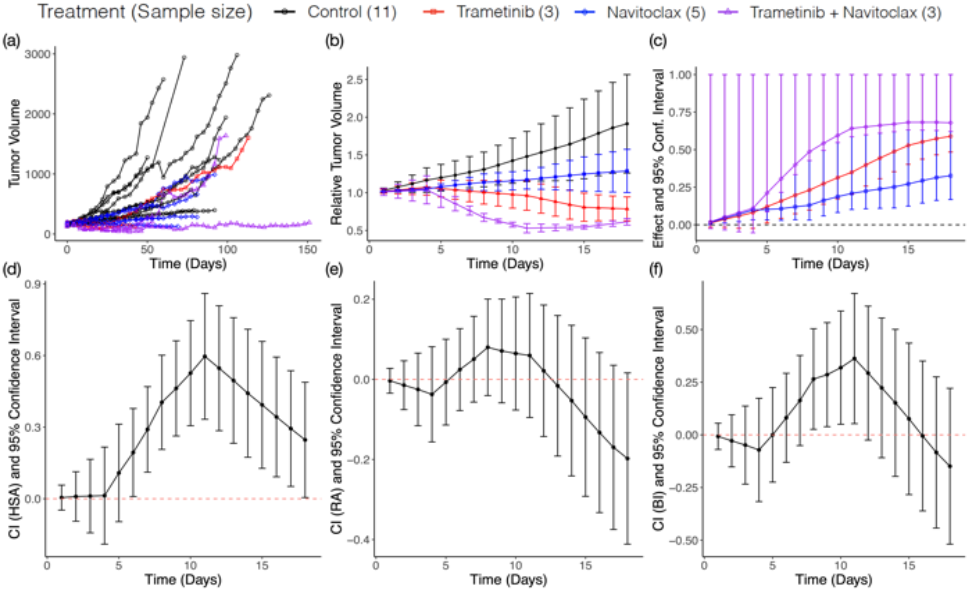
Effect and Combination Indices for trametinib plus navitoclax. (a) Profile plot of tumor volume. (b) Profile plot of relative tumor volume. Y-axis shows the mean ± standard error of relative tumor volume within each treatment group. (c) The drug effect for each treatment group relative to control group is quantified by tumor growth inhibition (TGI). The vertical line indicates the one-sided 95% confidence interval. (d)-(f) The joint action of combination drug under each reference model (HSA, RA, and BI) is assessed using a combination index. The vertical line indicates the two-sided 95% confidence interval.

### 3.3 Molecular Biomarkers Associated with CIs

Although KRT232 showed no synergistic signal in combination with navitoclax and trametinib (Table S5), we systematically investigated pathway-level signatures of the combination action in a framework of pharmacogenomic analysis utilizing multiple PDX experiments performed for the combinations. We conducted gene set variation analysis (GSVA) (Hänzelmann, et al., 2013) based on C2 collection of curated biological pathways as provided by the Broad Institute’s collection (Liberzon, et al., 2015). The pathway enrichment score (ES) by GSVA provides single-number summaries of pathway activity for each sample and each pathway. The PDX models were profiled for their expression of 15,732 genes using RNA-seq after filtering out genes whose 75% percentile was less than 20. Pathways with less than 10 genes were excluded, which resulted in 5164 pathways in total. We selected PDX models that had RNA-seq data available and fulfilled the Bliss assumption (0<*δ*_*g*_<1) across all time points. We came up with seven and nine PDXs in KRT232 plus navitoclax and KRT232 plus trametinib (Section S2).

We performed distance correlation tests (Székely and Rizzo, 2013) for detecting linear/nonlinear associations between pathways and the combination indices (Section S2). In combination treatment KRT232 plus navitoclax in NSCLC, controlling FDR at 0.1, resulting in 136, 150, and 145 pathways were significantly associated with HSA, RA, and BI, respectively, with 118 intersecting pathways across the three reference models (Figure 4a). Heatmap of those pathways associated with at least one of the *CIs* shows clear pattern of two clusters of PDX models (Figure 4b). The top significant pathways include those related to the therapeutic target or the prognosis in NSCLC (Figure 4c and Data S1). For example, the p53 pathway interplays with MDM2 inhibitor KRT-232 to suppress tumor cell growth (Rew and Sun, 2014; Sun, et al., 2014; Zhang, et al., 2020). Moreover, IRAK and HIF-1 pathways are associated with the development of tumor in NSCLC (Kurtipek, et al., 2016; Shimoda and Semenza, 2011; Zhang, et al., 2014).

**Figure 4:**
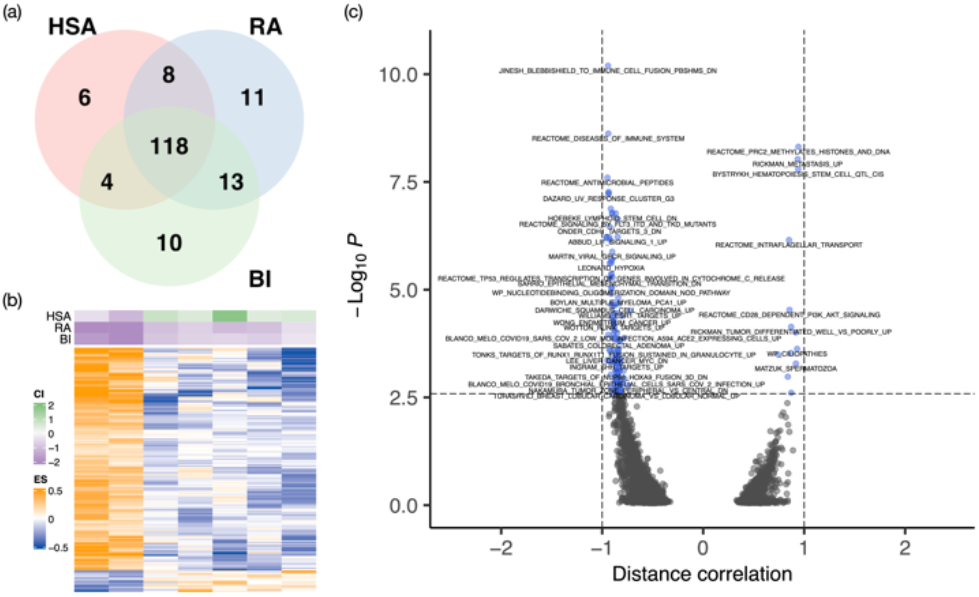
Differentially activated pathways at FDR of 0.1 for KRT232 plus navitoclax. (a) Venn diagram showing distribution of significantly expressed pathways. The figure illustrates the number of statistically significantly expressed pathways associated with each *gCI* (b) Heatmap of baseline pathway enrichment score from GSVA with the row annotation to be *gCI*. Each column represents a PDX model, and each row represents a gene whose enrichment score was significantly correlated with one or more *gCI* using the distance correlation test. (c) Volcano plot showing that p-value versus distance correlation for BI (the direction of the correlation is obtained by Spearman correlation). Blue dots represent significant differential pathways and black dots represent insignificant pathways.

Similar analysis was performed in combination treatment KRT232 plus trametinib. Controlling FDR at 0.1, resulting in 27, 12, and 13 pathways were significantly associated with HSA, RA, and BI, respectively, with seven intersecting pathways across the three reference models (Figure S3a and Data S1). Heatmap of those 32 pathways also shows clear pattern of two clusters of PDXs (Figure S3b). The seven intersecting pathways include those related to the therapeutic target in NSCLC. For example, several members in NFAT gene family differentially expressed in tumor vs. normal cells (Chen, et al., 2011), and FGFR3 is a potential therapeutic target in NSCLC (Chandrani, et al., 2017; Shinmura, et al., 2014).

## 4 Discussion

In this article, we have proposed a comprehensive statistical framework to quantify the joint action of two drugs in standard in vivo combination experiments with fixed doses, where the dose-response models such as the Chou-Talalay method (Chou and Talalay, 1983; Chou and Talalay, 1984) and Isobologram (Greco, et al., 1995) are not applicable. Our framework is generally applicable to tumor xenograft designs, including PDX experiments that is a newly designed novel model system for drug development and individualized treatment. The usual practice to decide combination effect in in vivo designs is based on p-values obtained from two-group comparisons of the combination group versus the control and monotherapy groups. However, the p-value subthresholding approach does not directly quantify the magnitude of the combination effect that is useful for further statistical modeling in pharmacogenomic setting where the driving molecular mechanisms of variable drug responses are systematically studied.

The combPDX web-application provides the visualization and analysis pipeline of the longitudinal tumor volume data at fixed dose levels, as well as power analysis tool to design in vivo combination experiments. Our framework is inspired by effect-based approaches that compare the effect from the combination of two drugs AB to the effects from its individual mono-drugs *A* and *B*, following the determination of efficacy of each treatment compared to the control. Various metrics have been suggested to summarize each individual growth curve to a value, e.g., the adjusted area under the tumor growth curve (aAUC) (Evrard, et al., 2020), which can be employed in our CI calibrations based on the fact that the aAUC is interpreted as the mean tumor volume across time.

There is no global consensus on defining drug synergism/antagonism in the field. Different reference models may lead to inconsistent statistical significance of combination effects based on different underlying assumptions. We have extensively studied the differences of these three models in the mathematical formulations. With fixed group means of C, A, B, and AB, the HSA model always provides the highest *CI* value, while RA has the lowest *CI* in that it provides the most conservative procedure to declare a synergistic effect (see Section S1.6). The CI derived under HSA is formulated similarly to t-tests that compare tumor volume data of combination therapy group vs. the better monotherapy group although the bootstrap procedure of HSA is more robust than the t-test. Both methods have the limitation of not utilizing tumor volume measurements from control group. Thus, the synergism declared from these methods should be considered as the minimal evidence and used for drugs which mono-therapeutic effects have been proved sufficiently in the field. In a counter example of drug antagonism (Figure S4) where the two drugs present a clear antagonistic pattern, HSA shows additivity because the tumor volume of the combination drug is close to that of trametinib, however, RA and BI, declared antagonism by adjusting the tumor volumes in control group (Table S5).

Using tumor volume data from NSCLC PDX experiments that evaluate two-drug combinations of KRT232, navitoclax, and trametinib, we have shown the utilities of calibrating *CI*s in finding underlying mechanisms of combination drug response based on gene expression data. We found that trametinib plus navitoclax had the synergistic effects in HSA and BI models, which is along with the currently undergoing clinical trials for the treatment of NSCLC (Kim and Giaccone, 2018). Moreover, the integrative analysis of KRT-232 plus navitoclax pharmacological data with gene expression data provided highly concordant pathway signatures across the three reference interaction models. These pathways included major driving mechanisms of the combination therapy in NSCLC.

In summary, combPDX represents an important step towards combination effect calibration for PDX models. It provides comprehensive data analysis and result visualization for in vivo combination drug testing. Coupled with molecular profile, combPDX facilitates the discovery of new biomarkers for combination therapy. Going forward, the knowledge of the biological mechanisms will add promise to the identification of the optimal personalized treatment.

## Data Availability Statements

Data analyzed in this article can be found online at doi:10.1016/j.canlet.2014.11.024.

## Funding

This work was supported by the National Institutes of Health 5U54CA224065-04 to J.W., B.F., F.M-B., J.R., and M.J.H. 2P50CA070907-21A1 to J.W., B.F., J.R., and M.J.H., 1R01CA244845-01A1 to M.J.H.

## Conflict of Interest

none declared.

## Notes

### Competing Interest Statement

The authors have declared no competing interest.

